# Functional roles of neural aPKCs in mouse brain development and survival

**DOI:** 10.1101/2024.05.22.595312

**Authors:** Aicha El Ellam, Emily J. Alberto, Maria E. Mercau, Dimitrius T. Pramio, Krishna M. Bhat, William M Philbrick, Deborah Schechtman, Carla V. Rothlin, Sourav Ghosh

## Abstract

Conserved protein complexes establish and maintain cell polarity. In turn, cell polarity is indispensable for fundamental developmental processes such as asymmetric division of stem cells and establishment of subcellular membrane polarization during cell differentiation. There are three well characterized polarity complexes. Atypical Protein Kinase C (aPKC) is a conserved constituent of the PAR complex that phosphorylates not only substrates within this complex, but also in the other polarity complexes. Outside of the polarity complex, aPKC regulates a myriad of cellular processes such as migration, metabolism, and survival. In mammals, two paralogs, *Prkci* and *Prkcz*, form the aPKC subfamily. Here, we characterized the expression of the *Prkci* and *Prkcz* paralogs, including a variant transcript of *Prkcz*, in the mouse brain and specific cells of the neural lineage. We generated a series of mice with individual and collective ablation of the two aPKC paralogs in neural stem cells, new-born neurons, astrocytes and NG2+ cells, as well as a mouse expressing kinase-inactive PRKCI in neural progenitors. We examine the effects of loss of aPKC paralogs or its kinase activity on gross brain development and organismal viability. Our results identify a critical window in neural progenitor differentiation wherein aPKC function is indispensable for neurodevelopment. Beyond this period, the ablation of even both aPKCs is characterized by a conspicuous absence of anticipated drastic effects. The genetic models developed might prove useful for further interrogating aPKC function in neurodevelopment and neuronal function or to reveal the role of polarity complex function in neurons, astrocytes and oligodendrocytes during stress, injuries or diseases.

## INTRODUCTION

Neural stem cells give rise to differentiated cell types in the brain including neurons, astrocytes and oligodendrocytes. This process involves polarization and asymmetric division of the stem cell. During neural differentiation in *Drosophila*, neuroblasts polarize to form distinct apical and basal cortical domains (1). Proteins such as Miranda, Prospero, Brat, Staufen and Numb form the basal complex, while a kinase, aPKC, and PAR proteins form the apical complex. *Drosophila* neuroblasts lacking aPKC displayed defects in polarity – PAR6 and LGL failed to localize apically, and Miranda was mislocalized to the apical cortex (2). In the absence of aPKC, *Drosophila* larva perishes at L2 stage (2). In mammals, there are three well-characterized protein complexes that function in cell polarity - the PAR/aPKC complex, SCRIB/LGL/DLG complex and CRBS/PALS/PATJ complex (3). Individual proteins within these complexes are typically encoded by multiple paralogs and some of the paralogs express multiple variant transcripts (3). For example, while *Drosophila* has a single aPKC gene, there are two aPKC paralogs in mammals – *Prkci* and *Prkcz*. The full-length transcripts of *Prkci* and *Prkcz* encode PRKCI also known as aPKCι, and PRKCZ also known as aPKCζ, respectively. These proteins are approximately 72% identical in amino acid sequence (4). Distinct from *Prkcz* transcript 1, another transcript from *Prkcz* (transcript 2) encodes for the kinase domain of aPKCζ *sans* the regulatory domain, and is called PKMζ (4). This unique transcript is expressed in neurons (5–7) as is *Prkci* (4, 6, 8). Reports on the role of mammalian aPKCs in neurodevelopment are inconsistent. For example, shRNA studies indicate that aPKCι (*Prkci*) and aPKCζ (*Prkcz*) were necessary for fundamental aspects of neurodevelopment such as precursor differentiation and migration. Haploinsufficiency of CREB-binding protein (CBP) and loss of its histone acetyltransferase activity results in Rubinstein-Taybi syndrome that includes intellectual disabilities (9, 10). *Cbp* ^+/-^ mice have behavioral deficits as well as display embryonic cortical precursor differentiation defects (11). aPKCζ (*Prkcz*) was demonstrated to phosphorylate and regulate CBP histone acetyltransferase activity and this phosphorylation of CBP was reported to be essential for precursor differentiation (11). Knock down of aPKCι (*Prkci*) and aPKCζ (*Prkcz*) in radial precursors by *in utero* electroporation revealed a role of aPKCι (*Prkci*) in maintaining radial precursors by inhibiting their transition to basal progenitors, while aPKCζ (*Prkcz*) functioned in radial precursor to neuron differentiation (12). By contrast to these *in vitro* and *in utero* shRNA knockdown experiments, genetic knockout experiments surprisingly indicated that *Prkcz* was entirely dispensable for neuronal differentiation and gross neurodevelopment (13, 14). Mice with neural-specific conditional knockout of *Prkci* (*Nestin* Cre^+^ *Prkci* ^f/f^) were also born in expected Mendelian numbers and were indistinguishable from wild-type (WT) littermates 5 days after birth (14). These mice died within 1 month after birth, nonetheless, the primary defect was demonstrated to be not a defect in precursor differentiation but rather a loss of adhesion junctions in the neuroepithelium (14).

aPKCs are multifunctional protein kinases with molecular roles beyond that in cell polarity, such as in downstream regulation of MAPK signaling, NF-κB pathway, p70S6 kinase signaling cascade and glucose transport (15, 16). Therefore, beyond development, these kinases can be expected to have a crucial role in survival and metabolic function of one or multiple cell types such as neurons, astrocytes and oligodendrocytes (17–19). Yet, the genetic ablation of *Prkci* in mature neurons using *Syn1*-cre or *Camk2a*-cre failed to reveal any apparent neuronal loss or degeneration (20). The characterization of paralog-specific expression of aPKCs in neural cell types such as astrocytes and oligodendrocytes remains incomplete. In the glial lineage, aPKCζ (*Prkcz*) was indicated to be necessary for GFAP-positive astrocytes and A2B5-positive oligodendrocyte precursor differentiation (11). Furthermore, aPKCζ (*Prkcz*) has been reported to play a role in astrocyte migration *in vitro* (21, 22). Astrocyte-specific genetic deletion of *Prkcz* and/or *Prkci* has not been tested, but the *Prkcz* ^-/-^ mouse phenotype suggests that PRKCZ function in astrocytes is dispensable for survival. During brain development, astrocyte precursors migrate and astrogenesis begins by E18 in the brain (23, 24). Defects in precursor differentiation and/or astrocyte migration therefore can be expected to lead to gross neurodevelopmental defects, yet the *Prkcz* ^-/-^ mouse lacks an egregious neurodevelopmental phenotype.

One possibility for a lack of gross phenotype could be redundancies or compensations. The overlapping expression of variant cell polarity proteins, at least in part, contributes to the lack of proper assessment of the specific or redundant functional role/s of these cell polarity kinases in mammalian neurodevelopment. It remains unknown whether redundancies and/or compensation account for the lack of expected effect with regards to aPKC gene targeting, whereas more acute gene silencing reveal critical functional roles (11, 12, 21, 22). aPKCι (*Prkci*) has been reported to compensate for the genetic targeting of PKMζ (25, 26). The combined knockout of both aPKC paralogs can reveal functional redundancy or compensation. For example, the loss of both paralogs in the intestine (*Villin*-cre ^+^ *Prkcz* ^f/f^ *Prkci* ^f/f^) was necessary and sufficient for tumorigenesis, while individual knockouts (*Villin*-cre ^+^ *Prkcz* ^f/f^ or *Villin*-cre ^+^ *Prkci* ^f/f^) were not (27). Compound knockout of *Prkci* and *Prkcz* in neurons or other neural cell types remains untested. Here we determine the endogenous expression of atypical PKC transcripts during neural development, as well as investigate their functional role in distinct neural cell types by genetic ablation of individual or both aPKCs.

aPKCs are primarily known for their protein kinase activity (28, 29). aPKCs phosphorylate substrates within the PAR complex such as PARD3, as well as in the other polarity complexes such as LGL (4, 8) and a variety of other substrates such as IRS, MAPK pathway, NF-κB pathway and JAK-STAT pathway (29–32). PKMζ is predicted to be a constitutively active protein kinase (33) since it lacks the atypical C1 domain that functions in autoinhibition (34, 35). Nonetheless, for PARD3 it was shown that aPKC can not only phosphorylate this substrate, but also remain in complex with PARD3 as an inactive kinase (36). The ability of aPKC to remain in complex with PARD3 in its inactive state is consistent with some kinases having a non-enzymatic, scaffolding functions (37). Kinase-independent function of aPKC has been shown in *Drosophila* follicle cell polarity (38). Similarly, kinase-independent function of mammalian aPKCs has also been demonstrated. aPKCζ and aPKCι interact with MEK5 in an EGF-inducible manner, which results in MEK5 activation (39). Importantly, kinase inactive aPKCζ (*Prkcz*) was sufficient for the induction of this EGF-dependent MEK5 activation (39). The function of aPKCι (*Prkci*) in *in vitro* neuronal differentiation of mammalian PC12 cells was indicated to be at least partly independent of aPKC kinase activity (40). Intriguingly, PARD6β amounts were reduced in *Camk2*-Cre^+^ *Prkci* ^f/f^ and *Syn*-Cre^+^ *Prkci* ^f/f^ mice (20), suggestive of a scaffolding function of aPKCι (*Prkci*) towards PARD6. Therefore, we also engineered a kinase-inactive aPKCι (*Prkci*) to explore whether its function in neurodevelopment depends on substrate phosphorylation or on kinase-independent scaffolding function.

## RESULTS

### *Prkci* and *Prkcz* transcript 2, but not *Prkcz* transcript 1, are expressed during forebrain development and neural differentiation

To investigate if distinct temporal expression may explain the functional difference between these paralogs, we examined the developmental expression of the mammalian aPKC paralogs during brain development. We used primers specific for *Prkci*, *Prkcz* transcript 1 or *Prkcz* transcript 2 to detect these transcripts in the embryonic brain and postnatal forebrain by qRT-PCR. Two of the three transcripts – *Prkci* and *Prkcz* transcript 2, but not *Prkcz* transcript 1 – were readily detectable at all the times assessed (**Figure 1A**). *Prkci* expression appeared not to change significantly from E12.5 to P60 (**Figure 1A**). The exception was the P21 time point, which showed a slight decrease when compared to E12.5. *Prkcz* transcript 2 did not change appreciably from E12.5 to E15.5. By contrast, from E17.5 to P60, *Prkcz* transcript 2 amounts increased significantly compared to that at E12.5 (**Figure 1A**). As internal controls for expected developmentally regulated expression of genes, we also measured the expression of the stem cell marker *Sox2* and *Nes*, which were expressed early between E12.5 to E17.5 and dramatically decreased thereafter as expected (**Supplementary Figure S1**). The neuronal marker *Dcx* peaked at E17.5 and gradually decreased by P10 (**Supplementary Figure S1**). The astrocyte marker *Gfap* was detectable at E17.5 and peaked at P5 (**Supplementary Figure S1**). *Gfap* expression declined after P5 to E17.5 levels by P60. The genes for mature oligodendrocytes *Mbp* and *Plp1* were also induced at specific times of development (**Supplementary Figure S1**). *Mbp* was detected at P10 and peaked at P21. The expression level declined at P30 and reached P10 level at P60. *Plp1* was detected at P21 and declined at P60 roughly by a half. The expression patterns of these genes were consistent with the temporally sequential process of neurogenesis followed by astrocyte differentiation and by the birth and maturation of oligodendrocytes. Surprisingly, we failed to detect *Prkcz* transcript 1 in the forebrain (**Figure 1A**), despite many publications having reported a functional role of the ∼70 kD PRKCZ protein in cells such as astrocytes (21, 22).

**Figure 1:**
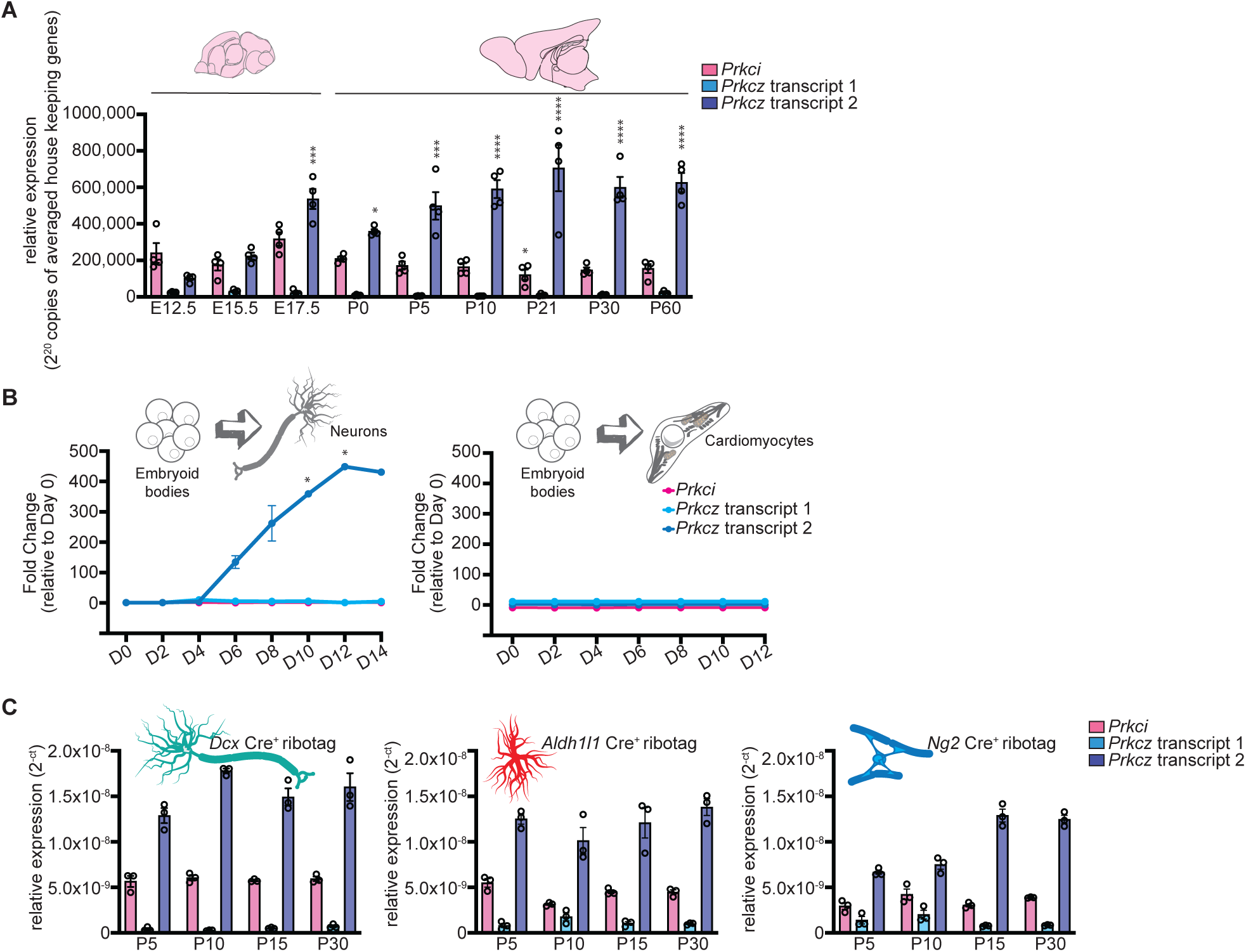
Expression of aPKC paralogs during forebrain development and neural differentiation. **A.** mRNA expression of *Prkci*, *Prkcz* transcript 1 and *Prkcz* transcript 2 in the brain of C57BL/6 mice at indicated embryonic (E) or postnatal day (P) days by qPCR. Data is mean ± SEM, n=4 mice. *p<0.05, ***p<0.001, ****p<0.0001 by one-way ANOVA, followed by Tukey’s test. **B.** mRNA expression of *Prkci*, *Prkcz* transcript 1 and *Prkcz* transcript 2 at the indicated time points during the differentiation of embryoid bodies into neurons (left) or cardiomyocytes (right). Data is mean ± SEM, n=3. *p<0.05 by one-way ANOVA, followed by Tukey’s test. **C.** Brains from *Dcx-cre*^+^ Ribotag ^Tg/+^, *Aldh1l1-cre*^+^ Ribotag ^Tg/+^, *Ng2-cre*^+^ Ribotag ^Tg/+^ mice were dissected at the indicated time-points and processed for Ribotag immunoprecipitation. mRNA expression of *Prkci*, *Prkcz* transcript 1 and *Prkcz* transcript 2 in each lineage and at the indicated postnatal (P) days was detected by qPCR. Data is mean ± SEM, n=3 mice.

*Prkcz* transcript 2 has been previously described to encode PKMζ (41). We observed that the expression of this transcript, as detected by our primers, was restricted to the brain tissue and excluded from other tissues examined, unlike *Prkcz* transcript 1 (**Supplementary Figure S2A**). Immunoprecipitation and immunoblotting experiments demonstrated that the presence of *Prkcz* transcript 2 indeed correlated with an ∼55 kD protein only in the brain (**Supplementary Figure S2B**). The forebrain lacked detectable ∼70 kD PRKCZ (**Supplementary Figure S2B**) consistent with our failure to detect *Prkcz* transcript 1 in this tissue (**Figure 1A, Supplementary Figure S2A**). Furthermore, when embryonic stem cells were differentiated *in vitro* along either a neuronal differentiation protocol or a cardiomyocyte differentiation protocol, *Prkcz* transcript 2 was induced exclusively along a neural differentiation lineage, and not during mesodermal (cardiomyocyte) differentiation (**Figure 1B**) suggesting it might be brain-restricted. Lineage-specific differentiation was validated by testing stem cell (*Oct4* and/or *Sox2* and/or *Nanog*), neuronal (*Msi1* and/or *Nes* and/or *Neurod1* and/or *Notch1* and/or *Tubb3*) and mesodermal (*Tbxt* and/or *Myh6*) markers (**Supplementary Figure S3**).

Next, we examined the expression of *Prkci*, *Prkcz* transcript 1 and *Prkcz* transcript 2 in neurons, astrocytes and oligodendrocytes at P5, P10, P15 and P30. We crossed lox-STOP-lox *Rpl22* exon 4-HA (ribotag) mice with *Dcx-cre*, *Aldh1l1-cre* and *Cspg4(Ng2)-cre*. Brain tissue was homogenized and RPL22^HA^-tagged polysomes were immunoprecipitated using anti-HA antibody. The ribosome-associated mRNAs were extracted and analyzed by qRT-PCR for *Prkci*, *Prkcz* transcript 1 and *Prkcz* transcript 2. Examination of enrichment of *Gfap*, *NeuN*, and *Olig2* and *Mbp*, respectively, confirmed the specificity of astrocytic, neuronal and oligodendrocyte-specific expression of ribosome-associated mRNAs (**Supplementary Figure S4**). Our results confirmed our whole brain expression analyses and indicated that *Prkcz* transcript 1 is not detected in any of the brain cell types investigated (**Figure 1C**). Interestingly, not only were *Prkci* and *Prkcz* transcript 2 expressed in neurons, these two transcripts were also expressed in astrocytes and oligodendrocytes (**Figure 1C**).

### Epigenetic changes correlate with brain-specific expression of *Prkcz* transcript 2

Transcription factors such as cAMP-responsive element binding protein (CREB) and CCAAT-enhanced binding protein (C/EBP) and CpG methylation has been implicated in the differential expression of *Prkcz* transcript 1 and *Prkcz* transcript 2 (41, 42). Furthermore, *Prkcz* transcript 1 and *Prkcz* transcript 2 promoters have been reported to respond differentially to HDAC inhibitors (43). To further examine the differential transcriptional regulation of *Prkcz* transcript 1 and *Prkcz* transcript 2, we examined CpG methylation and histone modifications around the putative start sites of these transcripts. We observed that the expression of *Prkcz* transcript 2 correlated with a brain-specific CpG-methylation pattern near its start site. Examination of CpG methylation patterns in several mouse tissues revealed a set of CpG islands were demethylated specifically in the forebrain and cerebellum but not in kidney or pancreas (**Figure 2A**). These tissue-specific demethylated CpG islands lie 3’ of the *Prkcz* transcript 1 start site between nucleotides +15615 to +15687, and likely correspond to the internal promoter for *Prkcz* transcript 2 expression. *Prkcz* transcript 2 expression also correlated with increased permissive H3K4 trimethylation and H3K9 acetylation at a nucleotide stretch +15092 to +16577 in the brain but not in kidney or pancreas (**Figure 2B**). Some enhanced CpG methylation was observed 5’ of *Prkcz* transcript 1 in the forebrain and cerebellum in the region of nucleotides +881 to +920 (**Figure 2A**), but there was clearly an increased enrichment of the repressive H3K27 trimethylation in the forebrain that correlated with the failure to detect *Prkcz* transcript 1 in this tissue (**Figure 2B**). Collectively, our results not only validate the expression of *Prkcz* transcript 2 in the brain, but also indicate that reduced tissue-specific CpG methylation and increased permissive histone modifications such as H3K4me3 and H3K9Ac correlate with the expression of this brain-specific transcript. The CpG methylation in tissues outside of the brain also suggests to brain-restriction of *Prkcz* transcript 2. By contrast, enhancement in CpG modifications at +881 to +920, and/or increased H3K27me3 in the forebrain prevent *Prkcz* transcript 1 expression in the brain.

**Figure 2:**
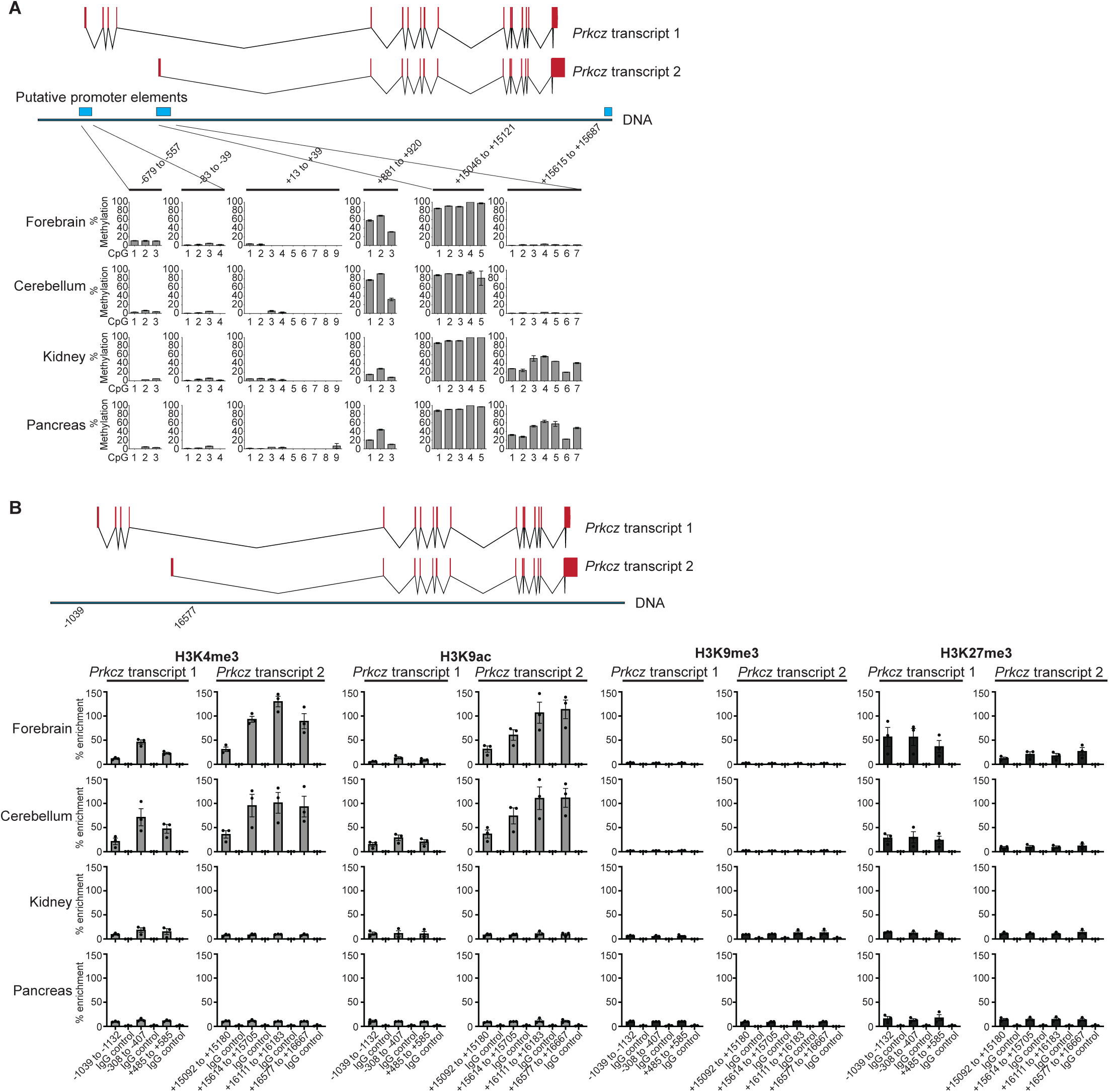
Differential CpG methylations and histone modifications correlate with tissue-specific expression of *Prkcz* alternative transcripts. **A.** Schematic of *Prkcz* transcripts 1 and 2, wherein exons are indicated in red bars, introns in black lines, Putative promoters in *Prkcz* locus is indicated by blue boxes and nucleotides of CpG regions are numbered. DNA was isolated from the indicated organs and tissues and % of CpG methylation of indicated regions was determined by bisulfite conversion and pyrosequencing. **B.** Schematic of *Prkcz* transcripts 1 and 2, wherein exons are indicated in red bars, introns in black lines, and relative positions of the genomic DNA regions investigated between 5’ from the *Prkcz* transcript 1 start site and 3’ from the alternative exon in *Prkcz* transcript 2 are labelled. Chromatin was isolated from the indicated organs and immunoprecipated with antibodies against the indicated modified histones. IgG was used as negative control. Enrichment of immunoprecipitated DNA, at the indicated positions, was determined by PCR. Data is mean ± SEM, n=3.

### PRKCI, including its kinase activity, is required in early neurodevelopment but is dispensable for viability after neural differentiation

It has been reported that *Nestin* Cre^+^ *Prkci* ^f/f^ mice died within 1 month after birth indicating an early role for PRKCI in neurodevelopment (14). We confirmed these findings, using an independently generated floxed *Prkci* mouse line (**Figure 3A**) crossed into *Nestin* Cre^+^. Excision of *Prkci* was confirmed by PCR (**Supplementary Figure S5A**). *Prkcz* gene was left intact (**Supplementary Figure S5B**). Immunoblotting confirmed a large reduction in PRKCI, but no changes in PKMζ signal (**Supplementary Figure S5C, D**). All *Nestin* Cre^+^ *Prkci* ^f/f^ died by 24 days after birth (**Figure 3B**). To test the cell-type specific function of PRKCI for survival at later stages of neurodevelopment following neural differentiation, we set up crosses to ablate *Prkci* in immature neurons (*Dcx* Cre^+^ *Prkci* ^f/f^ mice), astrocytes (*Gfap* Cre^+^ *Prkci* ^f/f^ mice) and oligodendrocytes (*Ng2* Cre^+^ *Prkci* ^f/f^ mice) and confirmed specific excision of *Prkci* but not *Prkcz* by PCR (**Supplementary Figure S5A, B**). Substantial reduction in PRKCI, but no changes in PKMζ, protein level were confirmed by immunoblotting in *Dcx* Cre^+^ *Prkci* ^f/f^ and *Gfap* Cre^+^ *Prkci* ^f/f^ mice. Only a modest reduction was observed in *Ng2* Cre^+^ *Prkci* ^f/f^ mice whole brains, perhaps due to fewer number of NG2^+^ cells compared to neurons and astrocytes (**Supplementary Figure S5C**). We calculated the expected and observed Mendelian ratios of F1 offspring genotypes for the following crosses: *Dcx* Cre^+^ *Prkci* ^f/+^ mice x *Prkci* ^f/f^ mice, *Gfap* Cre^+^ *Prkci* ^f/+^ mice x *Prkci* ^f/f^ mice and *Ng2* Cre^+^ *Prkci* ^f/+^ mice x *Prkci* ^f/f^ mice. There were no statistically significant differences between observed and expected Mendelian ratios of the genotypes at P24 in any of the above crosses (**Figure 3C**). The pups were phenotypically indistinguishable from wild-type, except *Ng2* Cre^+^ *Prkci* ^f/f^ mice, which were born smaller but caught up with wild-type littermates after 2 months (**Figure 3D**). The *Ng2* Cre^+^ *Prkci* ^f/f^ mice also had less hair, consistent with previously described expression of NG2 in hair cells (44, 45) and the role of PRKCI in hair follicle stem cells (46, 47). As previously reported, *Nestin* Cre^+^ *Prkci* ^f/f^ mice were significantly smaller and showed external signs of hydrocephaly (**Figure 3D**). We dissected the brain from these genotypes at P21 and performed histological analyses. These experiments revealed that there were no gross differences in the histology of the brain sections between wild type (WT) mice and *Dcx* Cre^+^ *Prkci* ^f/f^ mice, *Gfap* Cre^+^ *Prkci* ^f/f^ mice and *Ng2* Cre^+^ *Prkci* ^f/f^ mice, while *Nestin* Cre^+^ *Prkci* ^f/f^ mice had significant histological changes consistent with that reported in a previous publication by Imai *et al*. (14) (**Figure 3E**). These results demonstrated that while *Prkci* function in neural stem cells was essential for survival (*Nestin* Cre^+^ *Prkci* ^f/f^ mice), surprisingly, this was not the case in neurons, astrocytes and oligodendrocyte lineages.

**Figure 3:**
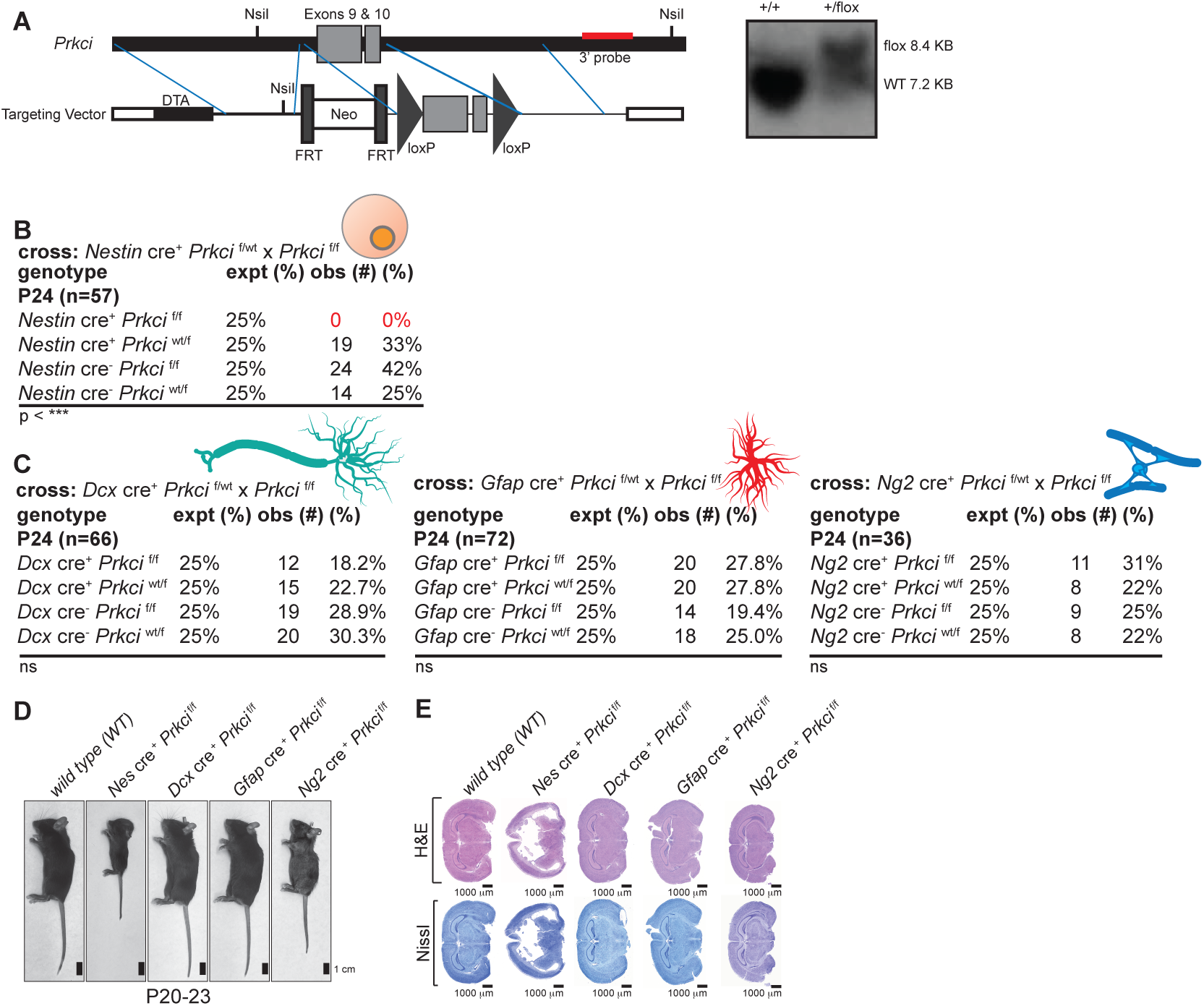
Genetic ablation of *Prkci* in neural stem cells, but not in neurons, astrocytes or oligodendrocytes, results in abnormal neurodevelopment and postnatal lethality. **A.** (left) Cloning strategy for the generation of *Prkci* ^f/f^ mice targeting the exons 9 and 10 and (right) identification of the *Prkci* floxed allele by southern blot making use of a 3’ probe indicated on the left. **B.** *Nestin* cre^+^ *Prkci* ^f/wt^ mice were crossed with *Prkci* ^f/f^ mice. Expected (expt) percentage (%) based on Mendelian ratio of offsprings for the indicated genotypes and observed (obs) number and percentage (%) are shown. ***p<0.001; Chi-squared test. **C.** *Dcx* cre^+^ *Prkci* ^f/wt^ (left) or *Gfap* cre^+^ *Prkci* ^f/wt^ (middle), or *Ng2* cre^+^ *Prkci* ^f/wt^ (right) mice were crossed with *Prkci* ^f/f^ mice. Expected (expt) percentage (%) based on Mendelian ratio of offsprings for the indicated genotypes and observed (obs) number and percentage (%) are shown. ns = non-significant; Chi-squared test. **D.** Representative photographs of mice of the indicated genotypes and age. **E.** Representative coronal sections of brains from mice of the indicated genotypes stained with H&E (top) or Nissl (bottom).

PRKCI has kinase activity towards several substrates, as well as kinase-independent functions (38–40). To test whether kinase activity is essential for the role of PRKCI in early neurodevelopment, we generated a kinase dead (KD) *Prkci* mouse model. First, we introduced point mutations in *Prkci* and tested their *in vitro* kinase activity and expression levels. Consistent with a previous report (48), mutating the ATP-coordinating lysine in PRKCI to arginine (K281R) did not show any reduction of kinase activity (**Supplementary Figure S6A**). However, mutating the DFG motif aspartic acid (D394) to alanine abolished kinase activity in *in vitro* assay (**Supplementary Figure S6A**). Neither of the amino acid substitutions altered protein expression (**Supplementary Figure S6B**). Next, we generated a *Prkci* KD mouse by introducing the D394A mutation in *Prkci* by CRISPR in C57BL/6 embryonic stem cells. The strategy for introducing base changes is shown in **Figure 4A**. Isolation of genomic DNA and sequencing of the targeting region identified a founder (#16) that was heterozygous for the introduced mutation (*Prkci* ^KD/WT^ mouse; **Figure 4B**). Heterozygocity at *Prkci* locus with one WT allele (*Prkci* ^KD/WT^) was viable and fertile, and the protein level of PRKCI in the brain was comparable to that with two WT alleles (*Prkci* ^WT/WT^; **Supplementary Figure S6C**). Crossing *Prkci* ^KD/WT^ to *Prkci* ^KD/WT^ mouse revealed that homozygous *Prkci* ^KD/KD^ mouse were never obtained, demonstrating that homozygous *Prkci* ^KD/KD^ is embryonic lethal (**Figure 4C**). Severe gross defects in embryonic development were apparent in the *Prkci* ^KD/KD^ embryos by E9.5 (**Figure 4D**). Similarly, gross defects in embryonic development were also exhibited by *Moex* Cre^+^ *Prkci* ^f/f^ mice, but not *Moex* Cre^+^ *Prkci* ^f/WT^ mice by E9.5 (**Figure 4E**). These results indicate that PRKCI function in embryonic development requires its kinase activity. Next, we investigated whether *Prkci* function in survival during neurodevelopment similarly required its kinase activity. Taking advantage of the fact that neither the heterozygous loss of *Prkci* or the presence of a *Prkci* ^KD^ allele affects survival, we generated a *Nestin* Cre^+^ *Prkci* ^f/WT^ x *Prkci* ^f/KD^ cross and counted the number of F1 offsprings. As expected, no *Nestin* Cre^+^ *Prkci* ^f/f^ mouse survived to P24 (**Figure 4F**). Identically, no surviving *Nestin* Cre^+^*Prkci* ^f/KD^ F1 offsprings were observed at P24 (**Figure 4F**). Both these genotypes displayed hydrocephaly (**Figure 3D, E and Figure 4G, H**). All other expected genotypes were observed and were grossly normal (**Figure 4F**). Examination of the brain tissue of *Nestin* Cre^+^ *Prkci* ^f/KD^ mice at P 20 revealed gross histological abnormalities (**Figure 4H**), similar to *Nestin* Cre^+^ *Prkci* ^f/f^ mice (**Figure 3E**). Thus, PRKCI function, including its kinase activity, is essential for viability early in development, including in neurodevelopment, while its expression during differentiation of neurons, astrocytes and oligodendrocytes is dispensable for mice viability.

**Figure 4:**
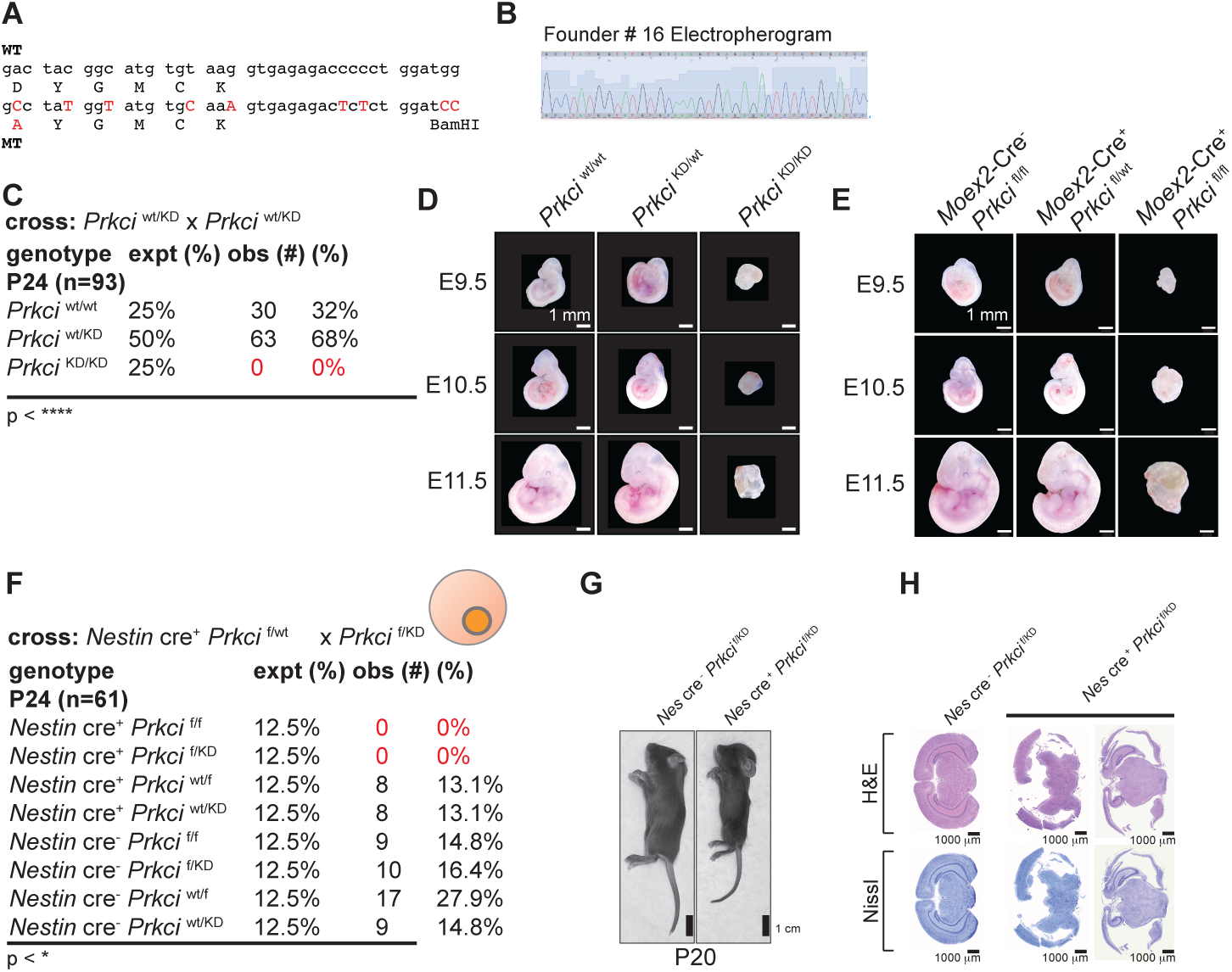
*Prkci* kinase activity is required for neurodevelopment. **A.** The DFG motif aspartic acid (D394) in *Prkci* was mutated to alanine to abolish its kinase activity. Changes in the genomic DNA introduced by CRISPR Cas9 are indicated in red. **B.** Electropherogram of a founder mouse, carrying the mutated sequence indicated in A. **C.** *Prkci* ^wt/KD^ mice were crossed with *Prkci* ^wt/KD^ mice. Expected (expt) percentage (%) based on Mendelian ratio of offsprings for the indicated genotypes and observed (obs) number and percentage (%) are shown. ***p<0.001; Chi-squared test. **D.** *Prkci* ^wt/KD^ mice were crossed with *Prkci* ^wt/KD^ mice. Representative images of embryos of the indicated genotypes and at the indicated embryonic (E) days. **E.** *Moex cre^+^ Prkci* ^f/wt^ mice were crossed with *Prkci* ^f/f^ mice. Images of embryos of the indicated genotypes and at the indicated embryonic (E) days. **F.** *Nestin* cre^+^ *Prkci* ^f/wt^ mice were crossed with *Prkci* ^f/KD^ mice. Expected (expt) percentage (%) based on Mendelian ratio of offsprings for the indicated genotypes and observed (obs) number and percentage (%) are shown. *p<0.05; Chi-squared test. **G.** Photographs of mice of the indicated genotypes at P20. **H.** Coronal sections of brains from mice of the indicated genotypes stained with H&E (top) or Nissl (bottom).

### PRKCZ requirement in early neural development is revealed in the absence of PRKCI but it cannot entirely substitute for PRKCI function

Given that *Prkci* and at least *Prkcz* transcript 2 are present throughout development in all post-natal differentiated neural cell types, we wanted to investigate if the paralogs function in survival during neurodevelopment. We noted that no effect on development and survival of mice has been reported even after germline ablation of *Prkcz* (13, 49–52). Presence of detectable *Prkcz* transcript 2 as early as E12.5 forestall completely disregarding a role of this transcript in neural development. Since the loss of *Prkci* causes death only around P24, it is possible that there might be some degree of functional redundancy between *Prkci* and *Prkcz* at very early neurodevelopmental time points. Albeit, in the presence of PRKCI the loss of *Prkcz* is entirely inconsequential for neurodevelopment. Therefore, to investigate if *Prkcz* loss has any effect on early neurodevelopment that affects viability, we generated a *Nestin* Cre^+^ *Prkci* ^f/+^ *Prkcz* ^-/-^ x *Prkci* ^f/f^ *Prkcz* ^-/-^ cross and examined the expected *versus* observed genotypes of the F1 offsprings at P0, P15 and P24 (**Figure 5A**). We also compared the percentages of observed and expected genotypes of *Nestin* Cre^+^ *Prkci* ^f/+^ x *Prkci* ^f/f^ cross at identical time points (**Figure 5B**). Chi-squared analyses revealed that there were no significant differences in the percentage of offsprings of each genotype observed relative to expected from either the *Nestin* Cre^+^ *Prkci* ^f/+^ *Prkcz* ^-/-^ x *Prkci* ^f/f^ *Prkcz* ^-/-^ cross or the *Nestin* Cre^+^ *Prkci* ^f/+^ x *Prkci* ^f/f^ cross at P0 (**Figure 5A, B**). Notably, *Nestin* Cre^+^ *Prkci* ^f/f^ *Prkcz* ^-/-^ and *Nestin* Cre^+^ *Prkci* ^f/f^ mice were born at the expected frequencies (**Figure 5A, B**). By contrast, at P15, there was a statistically significant difference between percentages of observed and expected genotypes in the *Nestin* Cre^+^ *Prkci* ^f/+^ *Prkcz* ^-/-^ x *Prkci* ^f/f^ *Prkcz* ^-/-^ cross but not in the *Nestin* Cre^+^ *Prkci* ^f/+^ x *Prkci* ^f/f^ cross (**Figure 5A, B**). Only one *Nestin* Cre^+^ *Prkci* ^f/f^ *Prkcz* ^-/-^ mouse out of 60 was alive at P15 and died immediately after (**Figure 5A**). *Nestin* Cre^+^ *Prkci* ^f/f^ genotype accounted for 15% of the F1 offsprings (10 out of 67 were alive) at P15 (**Figure 5B**). Albeit that this observed number of mice did not significantly differ from the expected numbers, there might be a slight trend towards reduced survival for this genotype at P15. Nonetheless, the reduction in survival dramatically increased in *Nestin* Cre^+^*Prkci* ^f/f^ *Prkcz* ^-/-^ at this age demonstrating that at P15 *Prkcz* can compensate for the loss of *Prkci*. This effect is lost by P24. At P24, both crosses showed a statistically significant difference in frequencies of observed genotype relative to expected genotypes (**Figure 5A, B**). These results indicate that both paralogs have at least partially overlapping roles in viability during neurodevelopment up to P15. While the loss of PRKCZ alone in the neural progenitors can be fully compensated by PRKCI, PRKCZ can also partially compensate for loss of PRKCI in the neural progenitors till this time point. The absence of both paralogs from neural stem cells leads to early lethality (P15). However, after this period, PRKCI function is essential. Absence of PRKCI alone from neural progenitors results in lethality and PRKCZ cannot compensate for the loss of PRKCI.

**Figure 5:**
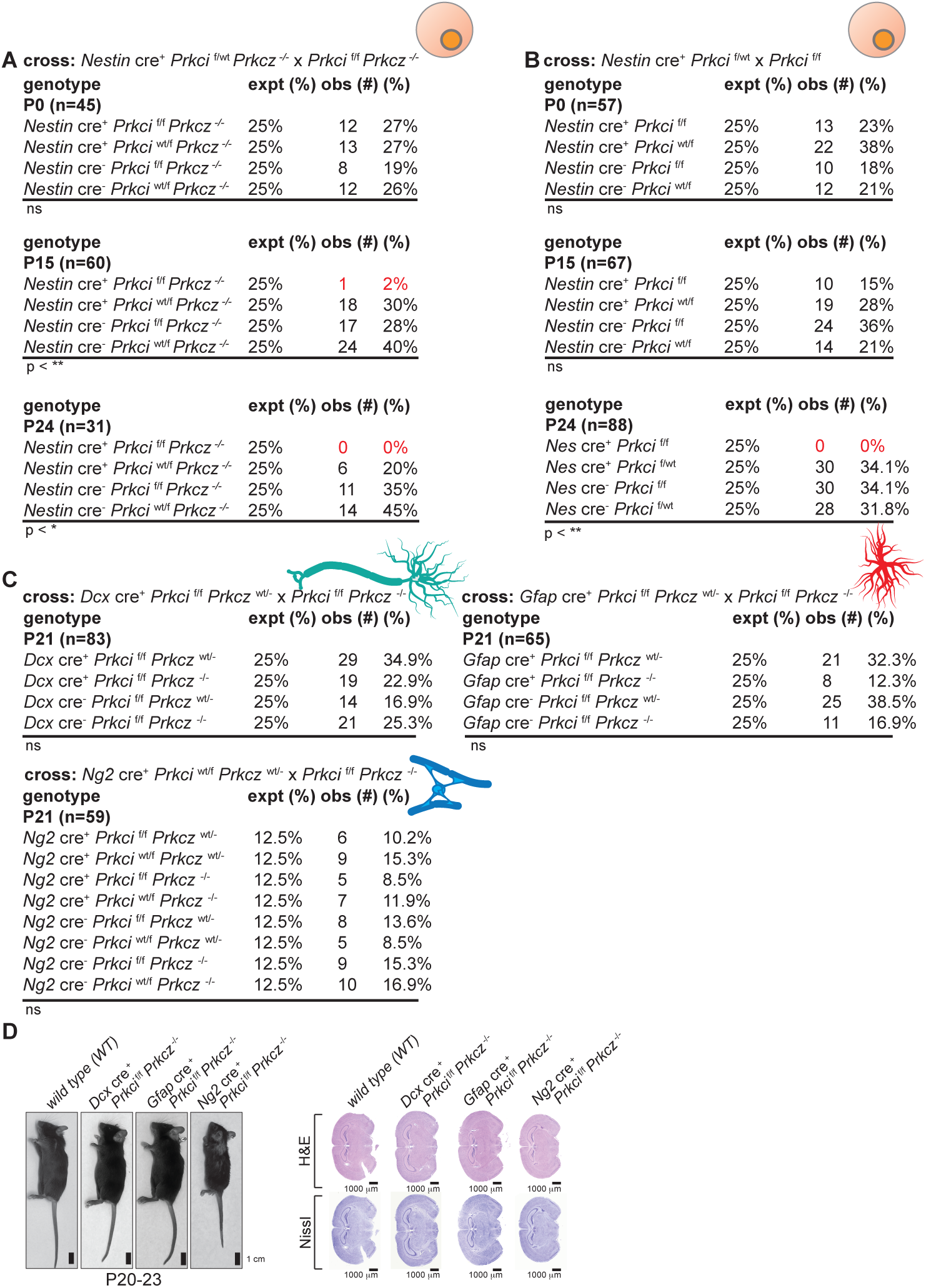
Genetic ablation of the paralog *Prkcz* accelerates postnatal lethality in *Nestin* cre^+^ *Prkci* ^f/f^ mice. **A.** *Nestin* cre^+^ *Prkci* ^f/wt^ *Prkcz* ^-/-^ mice were crossed with *Prkci* ^f/f^ *Prkcz* ^-/-^ mice. Expected (expt) percentage (%) based on Mendelian ratio of offsprings for the indicated genotypes and observed (obs) number and percentage (%) at P0, P15 or P24 are shown. *p<0.05, **p<0.01, ns=non-significant; Chi-squared test. **B.** *Nestin* cre^+^ *Prkci* ^f/wt^ mice were crossed with *Prkci* ^f/f^ mice. Expected (expt) percentage (%) based on Mendelian ratio of offsprings for the indicated genotypes and observed (obs) number and percentage (%) at P0, P15 or P24 are shown. **p<0.01, ns=non-significant; Chi-squared test. **C.** Dcx cre^+^ *Prkci* ^f/f^ *Prkcz* ^wt/-^ (top left), Gfap cre^+^ *Prkci* ^f/f^ *Prkcz* ^wt/-^ (top right) or Ng2 cre^+^ *Prkci* ^wt/f^ *Prkcz* ^wt/-^ (bottom left) mice were crossed with *Prkci* ^f/f^ *Prkcz* ^-/-^ mice. Expected (expt) percentage (%) based on Mendelian ratio of offsprings for the indicated genotypes and observed (obs) number and percentage (%) at P0, P15 or P24 are shown. ns=non-significant; Chi-squared test. **D.** (Left) Photographs of mice of the indicated genotypes and age. (Right) coronal sections of brains from mice of the indicated genotypes stained with H&E (top) or Nissl (bottom).

It is possible that *Prkci* function is only required at the neural stem cell stage and not once differentiation has occurred. *Prkcz* is entirely dispensable in this scenario. Alternatively, similar to early development, *Prkcz* transcript 2 is capable of a degree of compensation for *Prkci* in differentiated neurons, astrocytes and/or oligodendrocytes. We set up crosses to generate *Dcx* Cre^+^ *Prkci* ^f/f^ *Prkcz* ^-/-^, *Gfap* Cre^+^ *Prkci* ^f/f^ *Prkcz* ^-/-^ and *Ng2* Cre^+^ *Prkci* ^f/f^ *Prkcz* ^-/-^ mice. Immunoblotting validated a substantial reduction in PRKCI in the whole brain for *Dcx* Cre^+^ *Prkci* ^f/f^ *Prkcz* ^-/-^ and *Gfap* Cre^+^ *Prkci* ^f/f^ *Prkcz* ^-/-^. As observed before with *Ng2* Cre^+^ *Prkci* ^f/f^ mice, PRKCI protein was still detectable in *Ng2* Cre^+^ *Prkci* ^f/f^ *Prkcz* ^-/-^ brains (**Supplementary Figure S7**). PRKCZ protein was undetectable in *Dcx* Cre^+^ *Prkci* ^f/f^ *Prkcz* ^-/-^, *Gfap* Cre^+^ *Prkci* ^f/f^ *Prkcz* ^-/-^ and *Ng2* Cre^+^ *Prkci* ^f/f^ *Prkcz* ^-/-^ mice (**Supplementary Figure S7**). We observed no statistically significant differences between expected Mendelian ratios of F1 genotypes and observed genotypes in the F1 progenies of *Dcx* Cre^+^ *Prkci* ^f/+^ *Prkcz* ^-/-^ x *Prkci* ^f/f^ *Prkcz* ^-/-^ or *Gfap* Cre^+^ *Prkci* ^f/+^ *Prkcz* ^-/-^ x *Prkci* ^f/f^ *Prkcz* ^-/-^ crosses (**Figure 5C**). Although we could obtain *Ng2* Cre^+^ *Prkci* ^f/+^ *Prkcz* ^-/-^ and *Prkci* ^f/f^ *Prkcz* ^-/-^ mice for *Ng2* Cre^+^ *Prkci* ^f/+^ *Prkcz* ^-/-^ x *Prkci* ^f/f^ *Prkcz* ^-/-^ crosses, we consistently encountered difficulties in breeding. Therefore, we changed our strategy and established *Ng2* Cre^+^ *Prkci* ^f/f^ *Prkcz* ^f/+^ x *Prkci* ^f/f^ *Prkcz* ^-/-^ crosses. This cross gave expected progenies according to expected Mendelian inheritance, demonstrating that the absence of both *Prkci* and *Prkcz* did not grossly impact survival during neurodevelopment. Irrespective of their genotypes, the pups were grossly normal and indistinguishable, with the exception of *Ng2* Cre^+^ *Prkci* ^f/f^ *Prkcz* ^-/-^ that were slightly smaller in size and displayed hairless patches (**Figure 5D**). Additionally, we examined histological preparations of brain tissue of *Dcx* Cre^+^ *Prkci* ^f/f^ *Prkcz* ^-/-^, *Gfap* Cre^+^ *Prkci* ^f/f^ *Prkcz* ^-/-^ and *Ng2* Cre^+^ *Prkci* ^f/f^ *Prkcz* ^-/-^ mice to confirm that there were no gross differences with WT brain at P24 (**Figure 5E**). *Ng2* Cre^+^*Prkci* ^f/+^ *Prkcz* ^-/-^ mice, however, may not breed well due to unknown defects and possible function of aPKCs in NG2^+^ cells remain to be fully investigated. Collectively, these results demonstrate that PRKCI and, in its absence, PKMζ are required prior to neural progenitor differentiation, but not afterwards, in either neurons, astrocytes or oligodendrocytes for gross development of the brain and survival during this period.

## DISCUSSION

The investigation of aPKC variants in the adult mammalian brain revealed differential expression (53), however, knowledge of the endogenous expression of aPKC paralogs and transcript variants in the mammalian brain during its development is lacking. Furthermore, the expression of aPKC variants in astrocytes and oligodendrocytes have not been studied in detail. The absence of information on time-resolved and cell type-specific expression of aPKCs has led to experiments frequently employing ectopic expression or inappropriate targeting of aPKC brain-expressed isoforms, as well as the use of non-specific reagents (54, 55). These have occasionally led to paradoxical results, such as the differences between the studies pointing to a role of *Prkcz* transcript 1 *i.e.* aPKCζ in neuronal and glial differentiation (11, 12) and astrocyte migration (22) and genetic targeting of *Prkcz* (13). Differences in gene knockout and *in utero* shRNA targeting in brain development are not unheard of, as exemplified by the case of targeting *Dcx* (56–62). Redundancies, such as in the case of DCX family of proteins (56), off-target effects (63) or perhaps the lack of compensatory redundancies in the context of acute *versus* developmental ablation of a gene can account for differences between studies (56). Notwithstanding, differences have mired the study of aPKCs in controversies. For example, decades of study on PKMζ in learning and memory (64, 65) appeared to stand debunked (51, 52, 66), and then made a comeback (25, 26). Here we systematically investigate the expression of the aPKC variants in mice neurodevelopment, and determine the variant-specific expression in neurons, astrocytes and oligodendrocytes. We find that the brain-specific *Prkcz* transcript 2 encoding PKMζ protein is not only present in neurons, but also in astrocytes and oligodendrocytes. The function, if any, of this CREB and CBP regulated transcript (42) in these non-neuronal cells remains incompletely understood. Although experiments wherein *Prkcz* transcript 1 or aPKCζ was ectopically expressed in the brain or in astrocytes for functional studies can be found in published literature, this variant was entirely absent in the brain cells examined. We also identified differences in epigenetic mechanisms that may contribute to the selective expression of *Prkcz* transcript 1 and 2 in the brain and organs such as kidney and pancreas.

Previous studies indicate that functional compensation or redundancies between *Prkci* and *Prkcz* may be context dependent. A study employing substitution of all except the first 28 amino acids of PRKCI with the corresponding sequence of PRKCZ at the *Prkci* locus reported rescue of PRKCI loss by the chimeric PRKCZ at E7.5-8.5, but not at E9.5 (67). Another study demonstrated that the loss of either paralog was not sufficient, and the simultaneous loss of both was necessary for intestinal tumorigenesis (27). With regards to addressing functional consequences or redundancies and/or overlapping and non-overlapping functions of these paralogs, we not only tested the cell-type specific knockout of *Prkci* in neural progenitors, immature neurons, astrocytes and oligodendrocytes individually, but also combined these with germline ablation of *Prkcz*. We validated that the deletion of *Prkci* in neural progenitors results in hydrocephalus and death by 1 month after birth. Surprisingly, the deletion of *Prkci* in neurons, astrocytes or oligodendrocytes did not compromise birth or post-natal survival. Previously, it was documented that the ablation of this gene in mature neurons failed to reveal gross phenotypes (20). *Dcx*-cre is expressed in immature neurons at ∼ E15.5, much earlier than *Syn1*-cre, which is expected to lead to the ablation of *Prkci* approximately after 4-6 months of age, and *Camk2a*-cre, which is expected to ablate *Prkci* after ∼ 6 months of age (20). While genetic deletion of *Prkci* in astrocytes and oligodendrocytes were not tested before, our results with *Dcx* Cre-mediated ablation adds to the body of work on the functions of *Prkci* in neurons *in vivo*. The loss of *Prkci* in oligodendrocyte lineage may present with phenotypes that, albeit not essential for survival, deserve further examination. We also find that *Prkcz* transcript 2 or PKMζ can compensate for the loss of *Prkci* up to P15 but not at P24. However, even the loss of both did not affect neurons, astrocytes or oligodendrocytes. Taken together, there appears to be a temporal window when these kinases act redundantly before PRKCI activity becomes critical. The molecular basis for only a window when *Prkcz* transcript 2 or PKMζ compensates for the loss of PRKCI with PRKCI function becoming critical subsequently remains unknown. It is tempting to conjecture that the ∼86% amino acid identity between these paralogs in their kinase domains (4) allows for the redundancy, while the lack of domains – such as the PARD6 binding PB1 domain in *Prkcz* transcript 2 or PKMζ – limits the redundancy. Following the critical period of PRKCI requirement, the function of aPKCs appears to be dispensable for survival during neurodevelopment. Finally, in light of reports of kinase-independent functions of aPKCs (36, 38, 39), we also generated mice with *Prkci* kinase-inactive alleles replacing the endogenous alleles to test the functional role of its enzymatic activity. Our experiments unambiguously demonstrate that PRKCI biochemical activity in the developing brain requires its kinase activity and that this protein does not act exclusively as a scaffold.

Given the critical role of aPKC in establishing and maintaining polarity, the implication of the dispensability of these kinases for viability and development – even the loss of both paralogs from the genome – as it relates to cell polarity *per se*, requires further consideration. The polarity complex involving aPKC-PARD3-PARD6 is important for asymmetric division. aPKC polarity complex is also important for cell migration (68, 69) and the maintenance of structural and functional anisotrophy (70). aPKCs, given their limited number of paralogs and variants, allows for a systematic approach of characterizing expression and generating compound knockouts to investigate the role of cell polarity complex in the development of mammalian tissue and organs. This presents an opportunity more suitable when compared to the more complex variants in the case of PARD families (3, 8). For example, PARD6 has many paralogs in the mammalian genome (3, 8). PARD3 has only two paralogs but significantly more transcripts (3, 8). Our studies with aPKC, as far as neurodevelopment is concerned, appear to indicate that the essential function of the cell polarity complex surprisingly appears restricted to a specific but narrow window. Asymmetric division of stem cells is undeniably a key process for producing daughter cells with distinct fates (71–74). Alternative mechanisms that specify unique fates for daughters by suppressing identical fates in neighboring cells, such as Numb, or alternative cell biological or biophysical mechanisms of specifying asymmetry (75–77) may provide redundancies to polarity complex-driven asymmetry. Albeit that the mouse models developed for these studies unexpectedly failed to reveal gross differences in brain development, these may still be useful tools for interrogating brain function, including analyses of learning and memory in the adult brain. To this effect, a knock-in serine to alanine substitution of the aPKC phosphorylation site on CBP revealed defective adult hippocampal neurogenesis only at 6 months of age, but not at 3 months of age and diminished fear and spatial memory in mature adult (78). These tools might also prove useful in the study of brain function under conditions of stress, damage, injuries and diseases.

## ACKNOWLEDGEMENTS

This work was supported by Yale Funds (C.V.R. and S.G.). D.S. was a recipient of a FAPESP, BPE fellowship # 2015/24046-5. The authors would like to thank Irina Maskaykina for technical contributions.

## MATERIALS AND METHODS

### Animals

*Prkci* ^fl/fl^ mice were generated in 129/C57BL6 mixed background by standard techniques using a targeting vector containing a neomycin selection cassette flanked by FRT sites. Flox recombination sites were introduced in introns 5′ and 3′ of Exon 10 by homologous recombination (Fig. S2A). After electroporation of the linearized targeting vector into embryonic stem (ES) cells, G418 resistant ES cells were screened for successful homologous recombination by Southern blot analyses. Heterozygous recombinant ES clones were identified and microinjected into blastocysts from C57BL/6 J mice to generate chimeras. Germline transmitted floxed heterozygotes were selected and the neomycin cassette was removed by crossing with mice carrying the FLP recombinase. *Prkcz* ^fl/fl^ mice were previously reported in Chibly *et al* (49).

Kinase dead *Prkci* (*Prkci* D395A) mouse model was generated by CRISPR-Cas9 methodology as described in (79). Briefly, a T7-sgRNA transcription template was prepared by PCR, incorporating the guide sequence (CAGGTTACAAGCCATCCAGG, antisense orientation) from the desired target region in the mouse *Prkci* gene (NCBI Gene ID: 18759) with a predicted cut site within intron 12, 30 bp downstream of the targeted aspartic acid. The T7-sgRNA PCR template was then used for *in vitro* transcription and purification with the MEGAshortscript T7 Transcription Kit and MEGAclear Transcription Clean-Up Kit, respectively (both from ThermoFisher Scientific). Cas9 mRNA (CleanCap, 5-methoxyuridine-modified) was purchased from TriLink Biotechnologies. Homology-directed repair (HDR) oligonucleotide (168 base, including 65 base homology flanks) was ordered from Integrated DNA Technologies (IDT) as a phosphorothioate-protected Ultramer. C57BL6/N strain mice were obtained from Charles River Laboratories and cytoplasmic microinjections of sgRNA (50 ng/μl), Cas9 mRNA (100 ng/μl) and oligonucleotide homology-directed repair template (100 ng/μl) into single-cell embryos at 0.5 d pc were performed by the Yale Immunobiology CRISPR Core. DNA lysates of tissue biopsies from potential founder pups were screened by PCR amplification followed by a diagnostic BamHI digestion and then confirmed by sequencing. All animal procedures were performed according to NIH guidelines and approved by the Yale IACUC.

Other mouse lines used in this study were B6J.129(Cg)-*Rpl22*^tm1.1Psam^/SjJ or Ribo-tag [strain number 029977; JAX], B6.Cg-Tg(*Nes*-cre)1Kln/J or *Nestin*-cre [strain number 003771; JAX], *Dcx*-cre [strain number C57BL/6J-Tg(DCX-cre)35MullMmmh; Ulrich Mueller], B6.Cg-Tg(*Gfap*-cre)73.12Mvs/J or *Gfap*-cre [strain number 012886; JAX], B6;FVB-Tg(*Aldh1l1*-cre)JD1884Htz/J [strain number 023748; JAX] and Tg (*Cspg4*-cre/Esr1*)BAkik/J or *Ng2*-cre [strain number 008538; JAX]. Mice were bred and housed in a pathogen-free facility under standard conditions (12:12 light:dark cycle, with unrestricted access to food and water) and handled in accordance with the guidelines set by the Institutional Animal Care and Use Committee at Yale University and Yale Animal Resources Center. All the mouse strains used were backcrossed over 10 generations to a C57BL/6 background. Both male and female mice were used in experiments. All efforts were made to minimize the pain and the number of animals used. For timed pregnancies, embryos were collected at indicated dates, wherein morning of vaginal plug was considered E0.5.

### Genotyping

For confirming excision of the targeted gene, genomic DNA was extracted from mouse brain tissue and PCR amplifications were carried out with TopTaq Master Mix Kit (QIAGEN). PCR reactions of 25 μl were performed with 2 μl genomic DNA, 0.2 μM primer pair, 2.5 μl 10x CoralLoad Concentrate, 12.5 μl 2x TopTaq Master Mix (TopTaq DNA polymerase, TopTaq PCR Buffer with 3 mM MgCl2 and 400 μM each dNTP). PCR products were examined by gel electrophoresis. For *Prkci* KD mice, genomic DNA was extracted from ear punches. PCR products were purified using the QIAquick PCR Purification Kit (Cat. No. 28104) according to the manufacturer’s protocol. Between 40 to 140 ng of purified DNA was combined with the forward primer (*Prkci*FN). The purified DNA was sequenced at the Keck DNA Sequencing Facility at Yale and sequencing results were analyzed using SnapGene Software. Primers used for confirming excision and for sequencing are listed in **Supplementary Table 1**.

### RNA isolation and real-time quantitative PCR

Adult mice or mice at various days of postnatal development were euthanized by CO2 inhalation. Mice at various stages of embryonic development were collected by euthanizing the dam and then decapitating the embryos with surgical scissors. Brains and other tissues were dissected immediately on ice. Equal amounts of tissue were obtained from cerebral cortices using a 4 mm punch biopsy tool, flash frozen in liquid nitrogen and stored at −80° C until further processing. Total mRNA from brain tissue was extracted using RNAeasy Mini Kit (Qiagen) or TRIzol (15596026; Life Technologies), as per manufacturer’s instructions. RNA quantification and purity were assessed using a Nanodrop spectrophotometer. 1 mg of total RNA was used for cDNA synthesis. Reverse transcription was carried out using iScript cDNA synthesis kit (Bio-Rad Laboratories) or qScript cDNA Supermix kit (95048-100; Quanta Biosciences) and Real-time quantitative PCR (RT-qPCR) reactions were carried out were performed in triplicate using CFX96 Touch Real-Time PCR Detection System (Bio-Rad Laboratories) or ABI 7500 Fast Real Time PCR System (ThermoFisher Scientific) with PowerUp SYBR Green PCR Master Mix. For comparison of aPKC expression across tissues, ∼30 mg of tissues were extracted using RNeasy Mini Kit (Qiagen). 1 μg of total RNA was used for cDNA synthesis using iScript™ cDNA Synthesis Kit (Bio-Rad Laboratories). Standard curves were run for all primer pairs to ensure high efficiencies (>97%; R2 > 0.98) and a single shoulder-free peak upon melt curve analysis. One or more of *Gapdh*, *Rpl29* and *Eef1a* levels were used for normalization. Relative abundance of transcripts was calculated according to Pfafl method (80). Primer sequences are provided in **Supplementary Table 1**.

### Ribotag-qPCR

For Ribotag experiments, lineage-specific HA-tagging of ribosomal subunits was achieved by crossing RiboTag mice (81) to either *Dcx-cre*, *Aldh1l1-cre* or *Ng2-cre* recombinase drivers. Mice were sacrificed at the indicated postnatal time point by CO2 inhalation. Brains were dissected immediately, and 4 mm biopsies from cerebral cortices were flash frozen in liquid nitrogen and stored at −80° C until further processing. Ribotag immunoprecipitation and RNA isolation were performed following previously published protocols (82). Briefly, 200 μl of Dynabeads were washed and incubated in citrate-phosphate buffer with 10 μl of anti-HA antibody for 45 min at 4° C on a rotating platform. Cerebral cortices of 2 mice per time point were pooled and homogenized in homogenization buffer containing cycloheximide and RNAse inhibitors at a concentration of 10% m/v. After centrifugation at 10000xg for 10 min, 10% of the sample was dissolved in RLT buffer and frozen for further processing of INPUT. The remaining sample was incubated with the antibody-coupled beads on a rotating platform at 4° C for 6 hours. After washing, the beads containing the immunoprecipitated (IP) samples were washed and incubated with RLT buffer for 10 minutes on a thermomixer (750 rpm) at room temperature. Supernatants were collected and RNA was obtained from the IP and INPUT samples using QIAGEN RNeasy kits (Qiagen). Reverse transcription was carried out using iScript cDNA synthesis kit (Bio-Rad Laboratories), and PCR reactions were carried out using CFX96 Touch Real-Time PCR Detection System (Bio-Rad Laboratories).

### Western Blotting

Brains and other organs were dissected, diced and homogenized in radio-immunoprecipitation assay (RIPA) buffer (20 mM Tris, pH 7.2, 170 mM NaCl, 1% Sodium Deoxycholate, 1% Triton X-100, 0.1% SDS, 5 mM EGTA, 5 mM EDTA) containing cOmplete™ Protease Inhibitor Cocktail (Roche) by sonication. Protein concentration was determined by spectrophotometry using Pierce™ BCA Protein Assay Kits (ThermoFisher Scientific). Equal amounts (40 μg) of total protein in Laemmli Buffer were subjected to electrophoresis on precast polyacrylamide gels and transferred to PVDF membranes (Bio-Rad Laboratories). List of primary antibodies is provided in **Supplementary Table 2**. For detection of aPKCζ and PKMζ protein across mouse tissues, aPKC was immunoprecipitated from ∼70 mg of each tissue (C57BL/6 mice) in weak RIPA buffer (150mM NaCl, 25mM HEPES pH 7.5, 0,25% Sodium deoxycholate, 1% NP-40 and 10% glycerol) using 2 μg of anti-aPKCζ/PKMζ/aPKCι antibody (#SC17781, Santa Cruz) and 15 μl of Pierce™ Protein A/G Magnetic beads (ThermoFisher Scientific). Immunoprecipitates were separated on SDS-PAGE and transferred for immunoblotting. Membranes were blocked and probed overnight with anti-aPKCζ/PKMζ antibody (#C24E6, Cell Signaling Technology). Following washes, secondary antibodies conjugated to near infrared fluorophores (Li-COR Biosciences Cat #926-32213 and Cat #926-68072) or HRP (ThermoFisher Scientific #A16110) were incubated with the membrane. Blots were detected or developed using Odyssey Imaging System or Pierce™ ECL Western Blotting Substrate system (ThermoFisher Scientific) after further washes.

### ES cell differentiation (neuronal and cardiomyocyte)

Mouse ES cells were grown in DMEM-FCS/KOSR media with ESgro/LIF. Embryoid bodies were generated using hanging drop method (83), dissociated and grown in ITSF (insulin, transferrin, selenium and fibronectin) media on gelatin-coated plates for mesodermal differentiation (84) or in ADFNK (advanced DMEM/F12 neurobasal media with knockout serum replacement) media with retinoic acid and smoothened agonist (SAG) for neuronal differentiation (85).

### DNA methylation analysis

Genomic DNA was extracted from tissues indicated (30 mg) using the DNeasy Blood & Tissue Kit (Qiagen), according to the manufacturer’s instructions. 1 mg of genomic DNA was treated with sodium bisulfite using EZ DNA Methylation-Gold Kit (Zymo Research, Irvine, CA) according to the manufacturer’s instructions. The samples were eluted in 40 μL of M-Elution Buffer and 2 μL (equivalent to 25 ng of bisulfite-modified DNA) were used for each PCR reaction. Both, bisulfite conversion and subsequent pyrosequencing and analysis were done at the Epigenomics Profiling Core, at The University of Texas MD Anderson Cancer Center. PCR primers for the *Prkcz* locus were designed using the Pyromark® Assay Design SW 2.0 software (Qiagen). In brief, a sequencing primer was identified within 1 to 5 base pairs near the CpG sites of interest, with an annealing temperature of 40± 5° C. After that, forward and reverse primers were identified upstream and downstream to the sequencing primer, with a target annealing temperature ranging from 50° C to 60° C and amplicon product size ranging from 100 bp to 250 bp. Optimal annealing temperature was tested using gradient PCR. High methylation (SssI-treated DNA), low methylation (WGA-amplified DNA) and no-DNA template were included in the PCR reaction as controls. PCR reactions were performed in a total volume of 20 μl, and the entire volume was used for each pyrosequencing reaction, as previously described (86). Briefly, PCR product purification was done with streptavidin-sepharose high-performance beads (GE Healthcare Life Sciences, Piscataway, NJ), and co-denaturation of the biotinylated PCR products and sequencing primer (3.6 pmol/reaction) was conducted following the PSQ96 sample preparation guide. Sequencing was performed on a PyroMark Q96 ID instrument with the PyroMark Gold Q96 Reagents (Qiagen) according to the manufacturer’s instructions. The degree of methylation for each individual CpG site was calculated using the PyroMark Q96 software (Biotage AB, Uppsala, Sweden). The average methylation of all sites and triplicates was reported for each sample. Primer sequences are listed in **Supplementary Table 1**.

### Chromatin Immunoprecipitation

Chromatin isolation from fixed animal tissues was performed according to Cotney and Noonan protocol (87). Briefly, 30 mg of pancreas and kidney, and 70 mg of forebrain and cerebellum of adult C57BL6 mice were used for manual disruption and formaldehyde crosslinking. Nuclei was isolated and the chromatin was sonicated in a Branson SFX250 Sonifier (Branson Ultrasonics, Danbury, CT, USA), using 10% amplitude, 5s pulse and 5s rest, for a total of 7 minutes and 30 seconds. 5 μg of histone variants antibodies or IgG control antibody (please see **Supplementary Table 2**) were incubated overnight with 25 μg of chromatin. 5% of total chromatin without the antibodies was used separately as input alone control. Purified chromatin and input were quantified using the Qubit dsDNA HS (High Sensitivity) Assay Kit (ThermoFisher Scientific), and equal amounts of DNA were added to each qPCR reaction. Primer sequences are listed in **Supplementary Table 1**.

### Brain dissection and Histology

Mice were euthanized using carbon dioxide inhalation, brains were collected and fixed in 4% paraformaldehyde (PFA) for 24 hours. Fixed brains were transferred to 70% ethanol and shipped to iHisto Inc. (Salem, MA, USA) for embedding formalin-fixed paraffin embedded (FFPE) tissue, sectioning, deparaffinization, rehydration and Nissl or hematoxylin and eosin (H&E) staining on sections. Rostral telencephalon coronal sections were scanned using bright-field scanning, with a basic resolution of 40x objective lens, optical magnification of 40X, and 0.26 μm/pixel per slide.

### Kinase Assay

LanthaScreen TR-FRET Kinase Assay (ThermoFisher Scientific) was used according to manufacturer’s instructions. Briefly, HEK293 were transfected 3xFLAG-PKCι. FLAG-tagged proteins were pulled down using anti-FLAG M2 magnetic beads (Sigma) and serially diluted with enzyme buffer before ATP and fluorescein-conjugated PKC substrate peptide (ThermoFisher Scientific) were added and incubated. Reactions were extinguished with EDTA, incubated with Terbium-pSer PKC substrate antibody (ThermoFisher Scientific) and TR-FRET fluorescent signal was read on a microplate reader (Tecan).

