## supplementary material for "Functional roles of neural aPKCs in mouse brain development and survival"

### SUPPLEMENTARY FIGURE S1

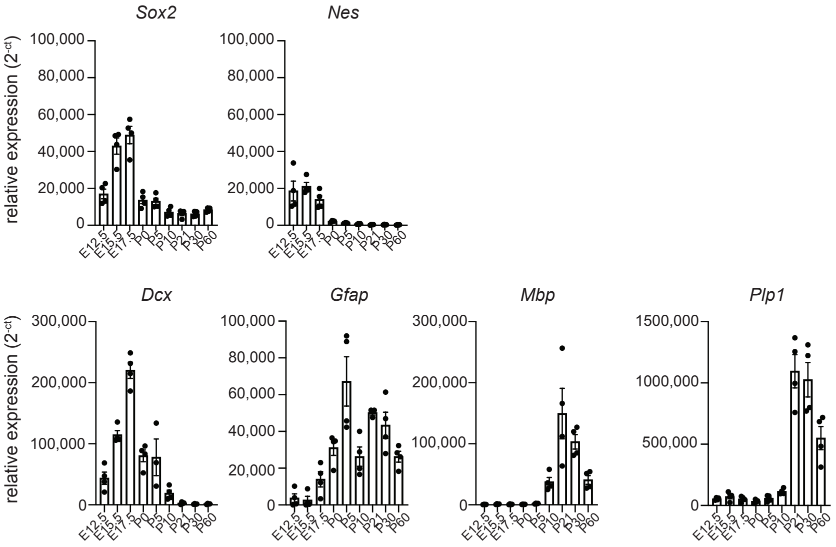

### SUPPLEMENTARY FIGURE S2

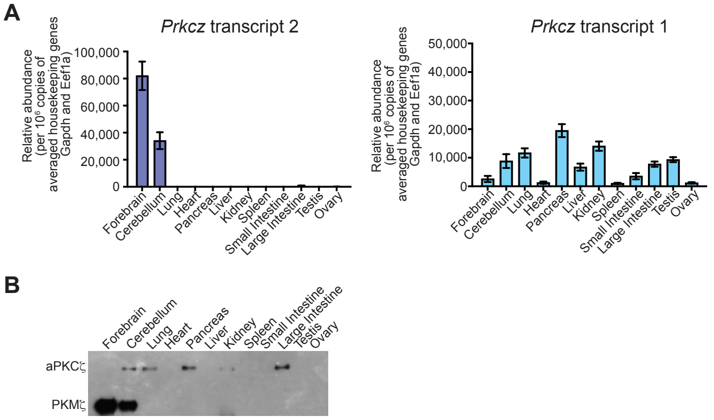

#### SUPPLEMENTARY FIGURE S3

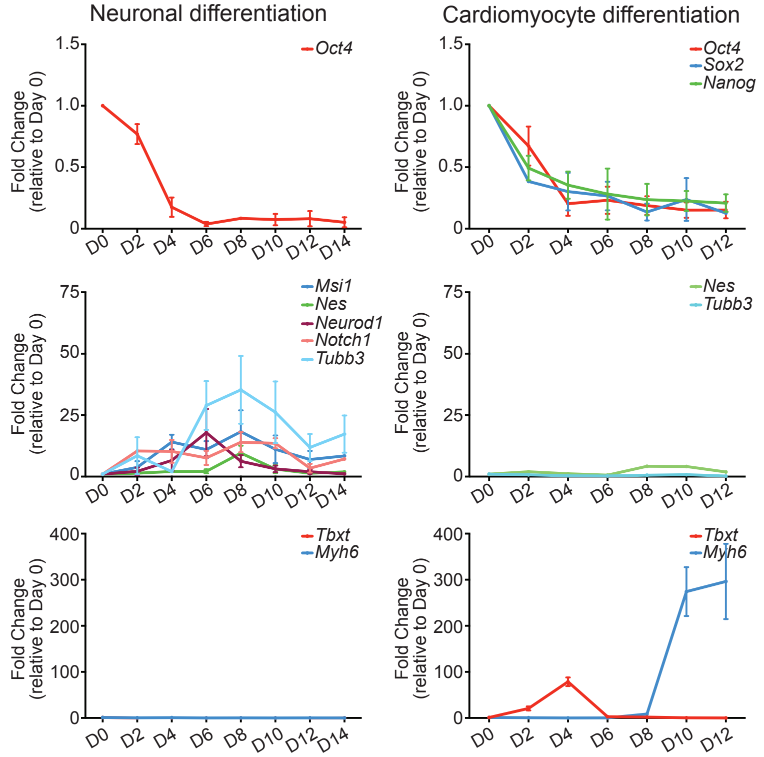

#### SUPPLEMENTARY FIGURE S4

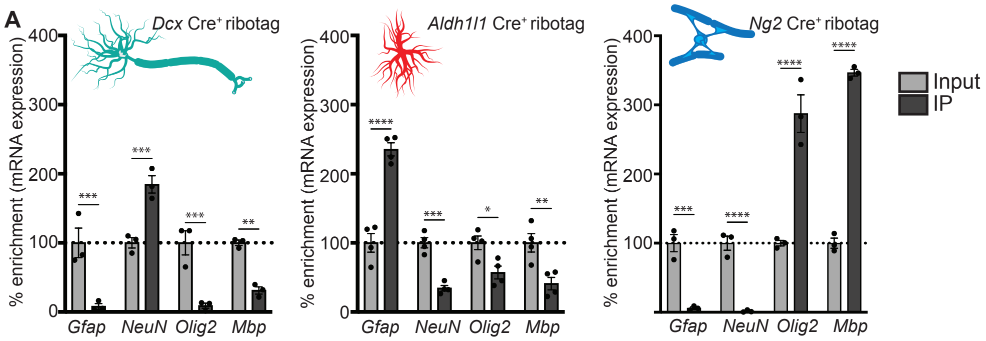

**SUPPLEMENTARY FIGURE S5**

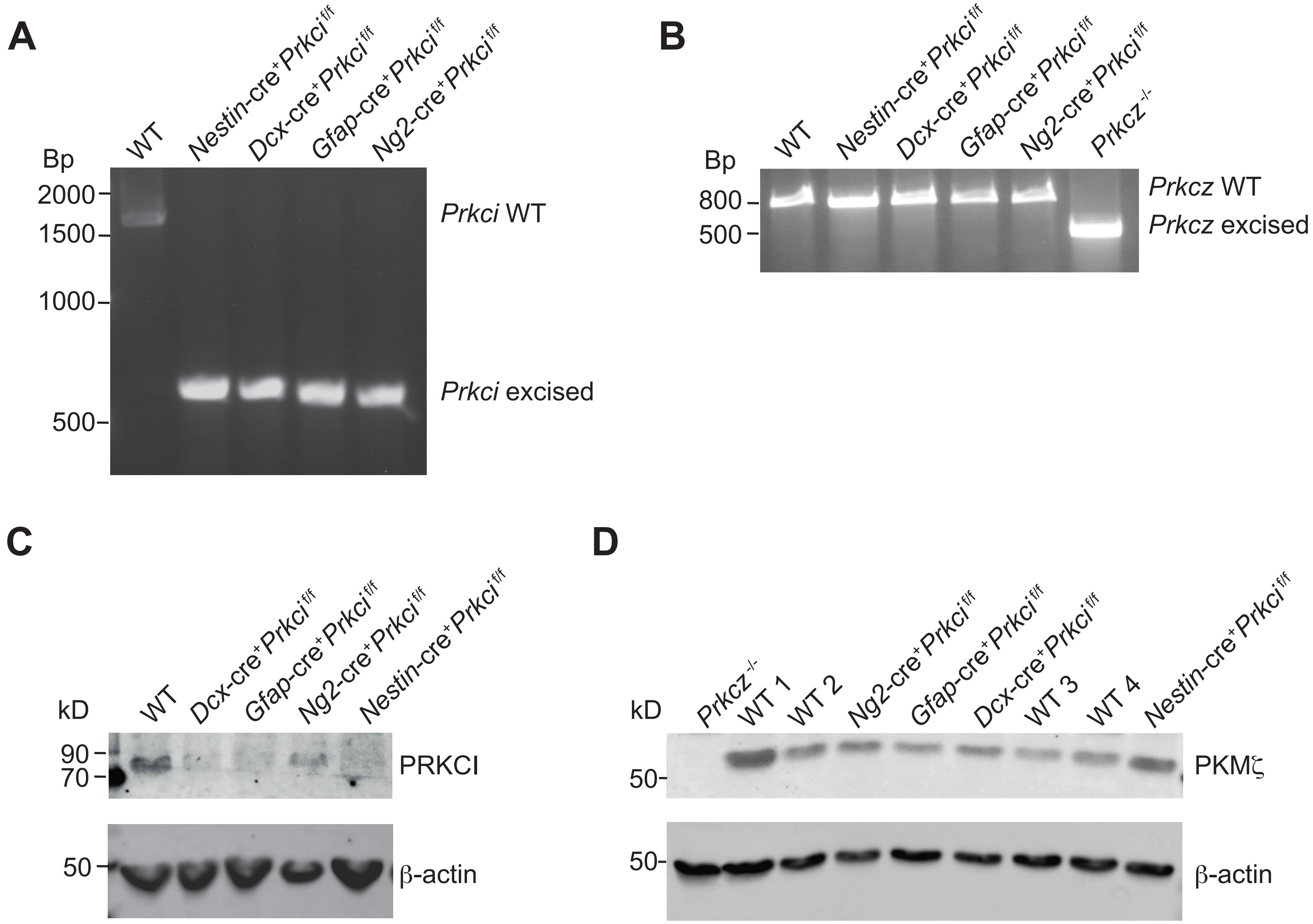

**SUPPLEMENTARY FIGURE S6**

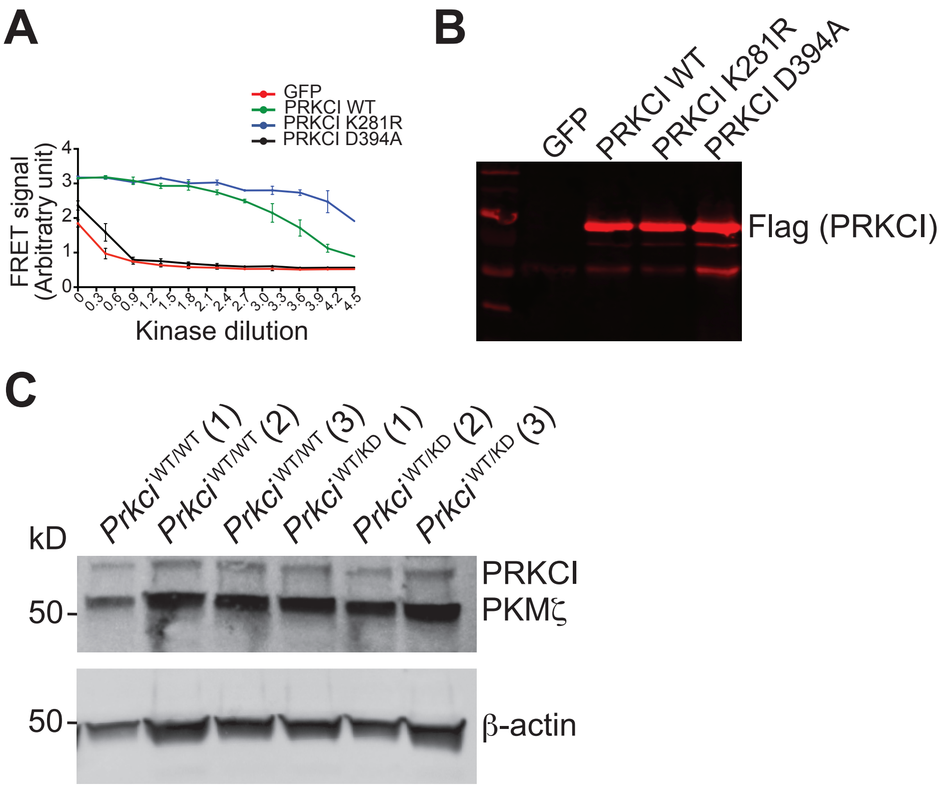

**SUPPLEMENTARY FIGURE S7**

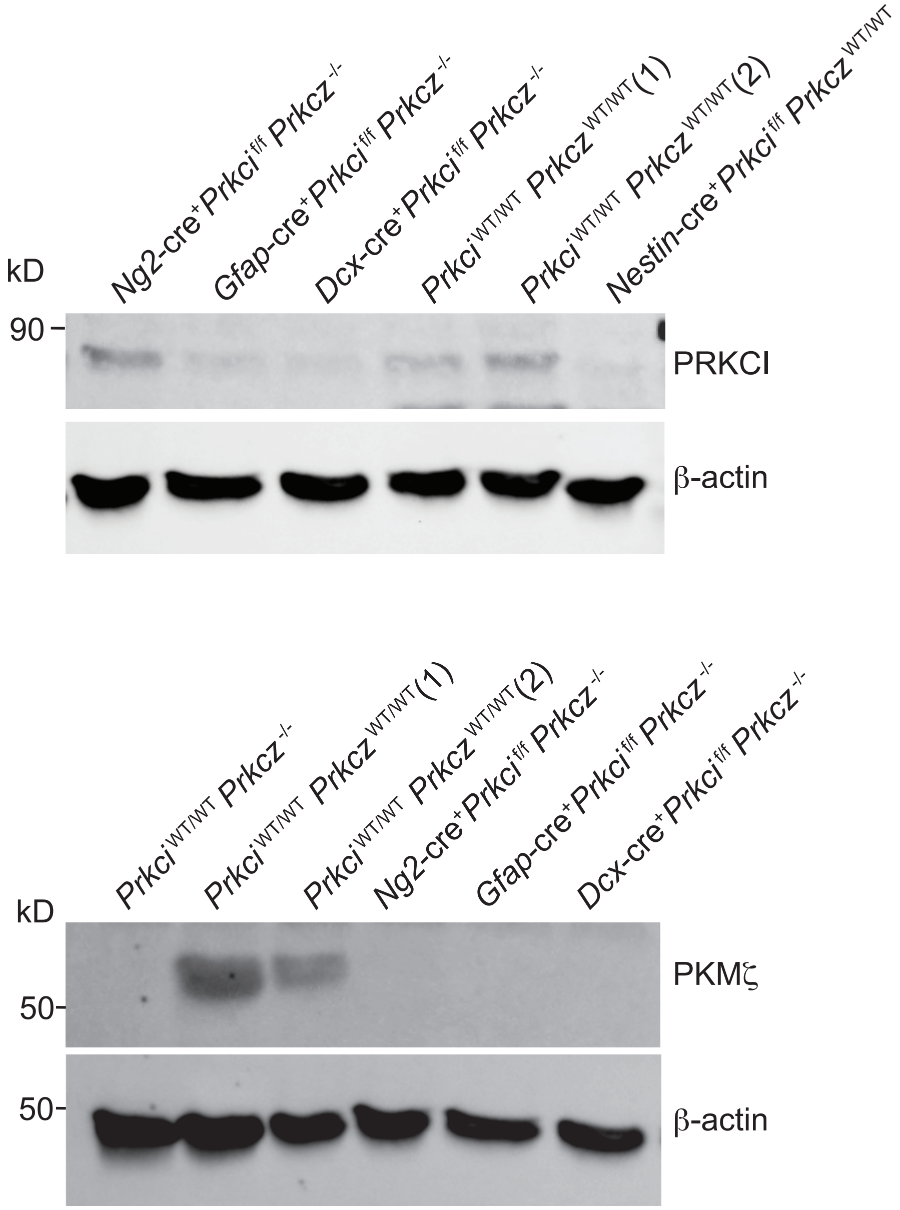

**Supplementary Figure Legends**

**Supplementary Figure S1: Gene expression of neural stem cell-, neuron-, astrocyte- and oligodendrocyte-associated genes during embryonic and postnatal development.** Brains were dissected at the indicated embryonic (E) or postnatal (P) time-points and indicated genes were detected by qPCR. Data is mean ± SEM, n=3-4, per time point.

**Supplementary Figure S2: Differential expression of *Prkcz* transcripts.**

**A**. Expression of *Prkcz* transcript 2 (left) and *Prkcz* transcript 1 (right) in the indicated organs, as detected by qPCR. Data is mean ± SEM, n=3.

**B**. Representative western blot for aPKCζ (encoded by *Prkcz* transcript 1) and PKMζ (encoded by *Prkcz* transcript 2) in the indicated organs.

**Supplementary Figure S3: Gene expression of stem cell-, neuronal- or cardiomyocyte-associated genes during embryoid body differentiation into neurons or cardiomyocytes.** mRNA expression of indicated genes at the indicated time points during the differentiation of embryoid bodies into neurons (left) or cardiomyocytes (right). Data is mean ± SEM, n=3.

**Supplementary Figure S4: Selected enrichment of genes associated with targeted populations by Ribotag approaches.** Brains from *Dcx-cre*$^{+}$ Ribotag $^{Tg/+}$, *Aldh1l1-cre*$^{+}$ Ribotag $^{Tg/+}$, *Ng2-cre*$^{+}$ Ribotag $^{Tg/+}$ mice were dissected and processed for Ribotag immunoprecipitation. mRNA expression for the indicated genes in the input and the IP fraction was detected by qPCR. Data is mean ± SEM, n=3 mice.

**Supplementary Figure S5: Specific ablation of PRKCI in conditional *Prkci* knockout mouse lines.**

**A**. Representative images of *Prkci* WT and excise alleles, as detected by PCR, in indicated genotypes.

**B**. Representative images of *Prkcz* WT and excise alleles, as detected by PCR, in indicated genotypes.

**C**. Representative western blot of PRKCI and β-actin loading control in indicated genotypes.

**D**. Representative western blot of PKMζ and β-actin loading control in indicated genotypes.

**Supplementary Figure S6:**

**A**. Lanthascreen™ TR-FRET kinase activity of Flag immunoprecipitates from HEK293 cells transfected with GFP alone (negative control) or PRKCI WT (positive control) and indicated mutants. Data is mean ± SEM, n=3.

**B**. Representative western blot of Flag-tagged PRKCI WT and indicated mutants from transfected HEK293 cells.

**C**. Representative western blot of PRKCI, PKMζ and β-actin loading control in brain lysate from mice with the indicated genotypes.

**Supplementary Figure S7: Genetic ablation of PRKCI and/or PRKCZ in tested mouse lines.**

**A**. Representative western blot of PRKCI and β-actin loading control in indicated genotypes.

**B**. Representative western blot of PKMζ and β-actin loading control in indicated genotypes.

**Supplementary Table 1**.

**qRT-PCR**

*Prkci*
CCATGTGTACCAGAGCGTCCT
TGTGGCCATTTGCACAATACA

*Prkcz transcript 1*
CAGGGACGAAGTGCTCATCA
CACGGCGGTAGATGGACTTG

*Prkcz transcript 2*
TGAGGAGGAGGCAGAGATGTG
GCAGGGTGGGTCTCCAGAT

*Sox2*
ATGGACAGCTACGCGCAC
CGAGCCGTTCATGTAGGTCTG

*Nes*
GCTGGAACAGAGATTGGAAGG
CCAGGATCTGAGCGATCTGAC

*Oct4*
GAAGCAGAAGAGGATCACCTTG
TTCTTAAGGCTGAGCTGCAAG

*Nanog*
ATGAAGTGCAAGCGGTGGCAGAAA
CCTGGTGGAGTCACAGAGTAGTTC

*Msil*
CCCAGGGTTCCAAGCCACGAC
GGGATAGCTGTGAGCTCGGGG

*NeuroD*
AAGGCAAGGTGTCCCGAGGC
AAGGCAAGGTGTCCCGAGGCT

*Notch*
ACACGGATGAGTGCGCCAGC
TGCCCGTTGAAGCCTTTGGGG

*Tubb3*
TAGACCCCAGCGGCAACTAT
GTTCCAGGTTCCAAGTCCACC

*Tbxt*
CATCGGAACAGCTCTCCAACCTAT
GTGGGCTGGCGTTATGACTCA

*Myh6*
GCAGAACAGTAAAATTGAGGACG
CGCAGCTTCTCCACCTTAG

*Dcx*
ACGACCAAGACGCAAATGGA
AATGACAGCGGCAGGTACAG

*Gfap*
AGAAAGGTTGAATCGCTGG
CGGCGATAGTCGTTAGCTTC

*Gapdh*
AGGTCGGTGTGAACGGATTTG
GGGGTCGTTGATGGCAACA

*Eef1a1*
CAACATCGTCGTAATCGGACA
GTCTAAGACCCAGGCGTACTT

*Rpl29*
CAAGTCCAAGAACCACACCAC
GCAAAGCGCATGTTCCTCAG

**Pyrosequencing**

ATTTTAGTGTTTGGTATTAAG
TCTACACCAATTAACTTC-biotinylated
Sequencing primer GGTATTAAGAAGAATTAGGT

TAGTTTATTTTGTATTTTGGTTAT
ACTAAATAAAAACCAAACCTAACCA-biotinylated
Sequencing primer GGATTGGTTTAGGTGG

GGTTTAGTTTTTATTTAGTGTG
TCTTAAAATCCCTCCTACTAAACATA-biotinylated
Sequencing primer ATTTAGTGTGTTTTTAGTTT

TTTAGGATAGGGATGATGG
ACCATACTAACCACTATCAT-biotinylated
Sequencing primer GGTTTTGGGGGTT

TTGAAGGGGTTGTGATTTGAATTTAGAG
CATCCACRACACAAAACTCACCTC-biotinylated
Sequencing primer TAAGTGTGGTTTGGTTAGTAG

ATGAGGTATTTTAAAGATT
TACCTCCTCCTCATATCT-biotinylated
Sequencing primer GTAGGTTTAGTATAGAGA

**Chromatin immunoprecipitation**

*Prkcz transcript 1*

GCACCAGAGGTTTTCACAATCTG
AGGTGTTGTTGCGCTCCTAG

AGGTGCGGGTTGTCCGATG
CGCACTTTGAGGAGTTGCTCAGC

GCTTCTGTCATGGCTATTAGCACTC
TGTCTGTGAACTCAGTGTGACACAC

*Prkcz transcript 2*

AAGGTAGGAGGTGGGCCTAAT
GAGGATCAAGCAGTCGGTCC

TCCCCTGGGAAGCTACGG
GAGACGCTTGTTTCCGTGTG

TTTCCTCCGACATCACCGC
ACAGATGTTCCAGCACGGAC

TGTGCACGTCAGAGCCTTTCG
AGCCTACCCTTGCCTTTCTGAG

**Genotyping primers (PCR and sequencing)**

*Prkci*

GGCTGAGGCACAAGGATAAC
CATGAGGTCCCCTCCATTTA

*Prkcz*

GAGAGCGAACCCTGAAAAGTG
AGTAAGCGGCAGCATAGGATA

Sequencing *Prkcz* kinase dead

TCTTCATGAGCGAGGGATAAT
GAAGGTGGCATGCAAATACTG

**Supplementary Table 2**.

Primary Antibodies:

Anti-MBP #808401 (BioLegend)
Anti-MOG #96457S (Cell Signaling Technology)
Anti-β-actin #3700S or Cat #8457S (Cell Signaling Technology)
Anti-aPKCζ/PKMζ/aPKCι #SC17781 or #SC216 (Santa Cruz)
Anti-aPKCζ/PKMζ #C24E6 (Cell Signaling Technology)
Anti-aPKCι #2998 (C83H11) (Cell Signaling Technology) or #SC11399 (Santa Cruz)
Anti-Flag M2 #F1804 (MilliporeSigma)
Anti-Histone H3 (acetyl K9) antibody #ab4441 (Abcam)
Anti-Histone H3 (tri methyl K4) antibody #ab8580 (Abcam)
Anti-Histone H3 (tri methyl K9) antibody #ab8898 (Abcam)
Anti-Histone H3 (tri methyl K27) antibody #ab6002 (Abcam)
